# *ExOrthist*: a tool to infer exon orthologies at any evolutionary distance

**DOI:** 10.1101/2021.02.22.432358

**Authors:** Yamile Márquez, Federica Mantica, Luca Cozzuto, Demian Burguera, Antonio Hermoso-Pulido, Julia Ponomarenko, Scott W. Roy, Manuel Irimia

## Abstract

Several bioinformatic tools have been developed for genome-wide identification of orthologous and paralogous genes among species. However, no existing tool allows the detection of orthologous/paralogous exons. Here, we present *ExOrthist*, a fully reproducible *Nextflow*-based software enabling to (i) infer exon homologs and orthogroups, (ii) visualize evolution of exon-intron structures, and (iii) assess conservation of alternative splicing patterns. *ExOrthist* not only evaluates exon sequence conservation but also considers the surrounding exon-intron context to derive genome-wide multi-species exon homologies at any evolutionary distance. We demonstrate its use in various evolutionary scenarios, from whole genome duplication to convergence of alternative splicing networks.

## Background

One of the most fascinating innovations of eukaryotic organisms was the evolution of gene structures composed of exons --pre-mRNA sequences joined and translated into proteins -- and interspersed introns --removed from the pre-mRNA through the process of splicing (Fig. 1A). Such innovation greatly increased the evolutionary potential of eukaryotic genomes.First of all, although rarely, changes in the exon-intron structure can modify the function of conserved genes (e.g. [1, 2]). More commonly, many eukaryotic genes acquire the ability to differentially combine their exons and introns into multiple isoforms through the process of alternative splicing. In particular, the inclusion or removal of an entire exon from an isoform (exon skipping) is the most common type of alternative splicing in metazoans and the main contributor to alternative splicing-driven proteome diversification [3]. Exon skipping has been shown to be fast-evolving [4–6], although some cassette exons are exceptionally conserved at the genomic and regulatory level [7]. Furthermore, in several cases, novel alternative splicing events have been linked to organismal innovations [8, 9]. Thus, exon-intron structure and alternative splicing patterns are two relevant intragenic features that have the potential to mediate functional diversification within conserved genes. However, how to link such features to function remains an open challenge [10, 11].

**Fig. 1:**
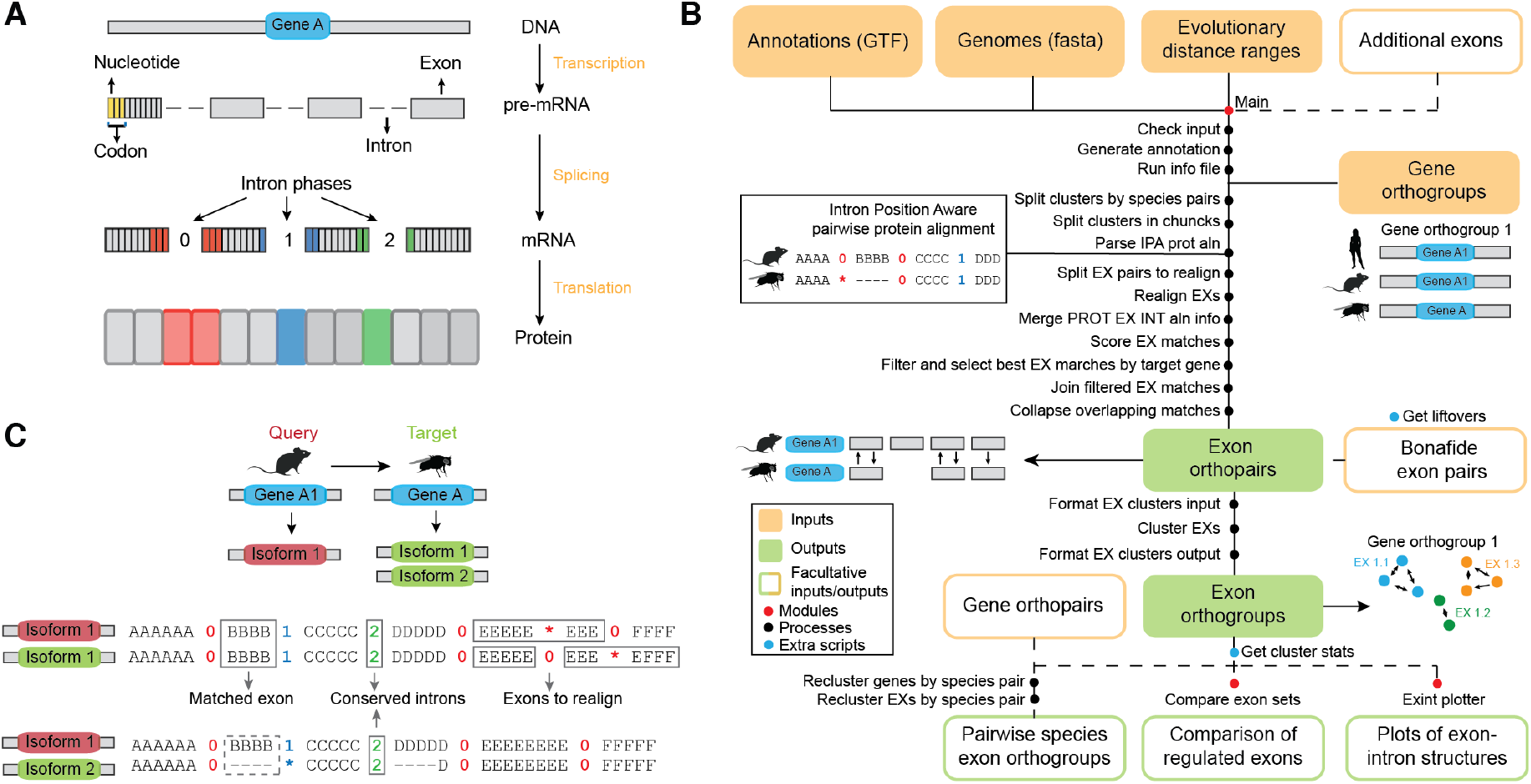
Molecular background and overview *of ExOrthist*. (**a**) Schematic representation of the exon-intron structure in eukaryotic protein-coding genes and of the molecular events necessary to generate a protein. The DNA is transcribed into pre-mRNA, from which introns are removed through the process of splicing. The exons are joined in the mature mRNA, where each adjacent group of three nucleotides (codon) encodes a different amino acid. Depending on the codon status at the exon-exon junctions, the removed introns are classified as being in phase 0 (complete codons on both sides of the junction) or phase 1 or 2 (split codons across the junction). The mature mRNA is then translated into a protein. (**b)** Overview of the *ExOrthist* pipeline, including inputs, outputs, processes, additional functionalities and modules. (**c**) Schematic representation of species pairwise IPA alignments generated by *ExOrthist*. Given a query and a target gene, *ExOrthist* aligns all non-redundant query gene protein isoforms against all target gene non-redundant protein isoforms. The figure highlights some common scenarios in IPA alignments: exon matches detected in only one of the aligned isoforms, introns with conserved phases and query exons matching multiple target exons (which will be specifically realigned).

Comparative genomics has been an important tool to link whole genes to functions by inferring gene presence/absence in different species through the identification of evolutionarily related gene orthologs (originated by a speciation event) and paralogs (derived from a duplication event). While orthologs usually maintain the ancestral function, paralogs are free to mutate and evolve new functions, as long as the ancestral role is preserved [12, 13]. Remarkably, although multiple bioinformatic tools have been developed to identify homology relationships at the gene level [14–18], and some are emerging to investigate the conservation of splicing patterns [19], no corresponding tool currently exists to infer exon homologies, particularly over deep evolutionary timescales. Nonetheless, even in the absence of general-purpose tools, a variety of studies have underscored the importance of this approach. First, in much the same manner as origins of novel genes have been linked to evolution of new molecular functions, *de novo* origins of alternatively spliced exons have been associated with evolution of novel gene functions [7, 8]. Second, in the same way as gene paralogs frequently evolve new functions, duplicated paralogous exons have been shown to provide novel functional properties to the host gene [20, 21]. Moreover, even in cases in which new functions do not emerge, mutually exclusive splicing of duplicated exon pairs can lead to the specialization of alternative transcripts for distinct pre-existing functions, mirroring the impact of whole gene duplication [22].

General tools for assigning and studying orthologous and paralogous exons are thus crucial to properly investigate the evolution of gene structures and alternative splicing. For a few model organisms, it is possible to confidently infer pairs of orthologs at short evolutionary distances (e.g. within mammals or within fruitflies) using *liftOver* [23]. Besides the restriction to short evolutionary distances, this approach relies on the availability of pre-computed UCSC pairwise chain alignments and can be applied only to one species pair at a time (i.e. multi-species orthogroups cannot be inferred). Moreover, *liftOver* derives exon orthopairs exclusively based on interpolation of the target exon sequence within whole genome alignments, which has two main limitations. First, it does not take into account the preservation of the two neighboring intron positions, whose ancestrality is a requisite for true exon homology. Second, it does not assess the conservation of the upstream and downstream exons, which helps to further support homology of the neighboring intron positions.

To overcome these limitations and fulfill the major need for a software for exon homology inference, we have developed *ExOrthist*, which uses Intron Position Aware (IPA) protein alignments to infer species pairwise exon homologs and multi-species orthogroups, with substantial flexibility to allow fine tuning according to evolutionary distance. *ExOrthist* has three modules. The *main* module is a *Nextflow*-based pipeline [24] that infers exon homologies using an innovative approach, namely evaluating three features: conservation of the neighboring intron positions, sequence and length conservation of the query exons, and sequence conservation of the upstream/downstream exons. Second, the *exint_plotter* module is also a *Nextflow*-based pipeline that allows the visualization of evolutionary conservation and of changes in exon-intron structure among gene homologs of interest. Third, the *compare_exon_sets* module assesses the conservation of alternative splicing patterns between pairs of species. Here, we provide the first description of *ExOrthist*. We first give an overview of the pipeline (schematized in Fig. 1B), describe how we selected the default conservation cut-offs for the three evolutionary distance ranges, and compare *ExOrthist* and *liftOver* performances on a set of closely related mammalian species. Then, we illustrate the use of *ExOrthist* to: (i) reconstruct paralogous relationships and asymmetric evolution after a whole genome duplication event in the *Xenopus* lineage; and (ii) compare *Nova*-regulated exons in mouse and fruitfly. All *ExOrthist* modules are freely available at https://github.com/biocorecrg/ExOrthist.

## Results

### Overview of *ExOrthist main* module

The *ExOrthist main* module infers exon homologies within a set of species for which annotation files (in GTF format) and genome files (in fasta format) are provided. Moreover, it is possible to provide additional non-annotated exons identified from RNA-seq data by any splicing quantifier (e.g. [25–29]). *ExOrthist* derives exon homologies at two different levels: exon orthopairs, representing pairs of homologous exons from two different species, and exon orthogroups, which include all the exons sharing a common ancestor (and thus both orthologs and paralogs) for multiple species. As such, orthopairs and orthogroups provide complementary information that is necessary for the complete reconstruction of exon evolutionary relationships. Since, by definition, exon orthopairs/orthogroups are derived from evolutionarily related genes, *ExOrthist* also requires as input a set of gene orthogroups for the species of interest (e.g. generated by *Orthofinder* [17], *Broccoli* [18] or similar tools). While *ExOrthist* is designed to infer genome-wide exon homologies, it can also be restricted to one/few genes of interest by providing only a subset of gene orthogroups as input. *ExOrthist* utilizes a set of conservation cut-offs (number of conserved intron positions, minimum exon sequence similarity, minimum exon length ratio, minimum global protein similarity; see below), which can be differentially fine-tuned for three evolutionary distance ranges (short, medium and long). Thus, the evolutionary range between each pair of considered species also has to be specified (see online README for further details on the input).

Once launched, *ExOrthist* extracts exon/intron-related information from the annotation files of all the species considered, and then it performs the subsequent steps for each pair of species. First, it generates IPA alignments for all non-redundant protein isoforms belonging to the same gene orthogroup for each species pair (species1-species2) (Fig. 1C). This all-vs-all isoform comparison is necessary because a selection of representative isoforms might leave some exon orthologs undetected simply because they are not included in any of the selected protein isoforms. Considering species1 as query and species2 as target (and vice-versa), *ExOrthist* then parses all pairwise isoform alignments. For each query exon, it first identifies the best matching exon in each target isoform based on the global protein alignment and sequence similarity. Next, it selects the best exon match for each target gene among all available isoforms. This selection is done based on a global score [0-1] given by the sum of five partial scores reflecting the conservation of different features of the exon-intron context: (1, 2) conservation of the immediately upstream and downstream intron positions and phases, (3) conservation of the query exon sequence and (4, 5) conservation of the immediately upstream and downstream exon sequences. Finally, for each best exon match per target gene, *ExOrthist* evaluates its potential homology with the query exon according to the conservation cut-offs provided (see below). If the query and target exons involved in this match pass all cut-offs, they are considered as homologs (i.e. an orthopair). Importantly, while the *ExOrthist* logic requires a query exon to match a unique exon in the target gene, each exon in the target gene can potentially be matched by multiple query exons: this setting captures cases of in-tandem exon duplication while preserving the information about which duplicated exon is more similar to the ancestral one. Furthermore, the user has the possibility to provide custom *bona fide* exon orthopairs, which will be directly incorporated into the generation of the exon orthogroups. This can be obtained from manual curation (e.g. through literature searches), or in a genome-wide manner using the *liftOver* tool. For this purpose, *ExOrthist* provides an additional script to perform *liftOver* between any pair of species for which UCSC *liftOver* files are available.

After deriving species pairwise exon homology relationships, *ExOrthist* infers multi-species exon orthogroups. The exon orthopairs for all the species pairs are joined and translated into a directed graph using the R *igraph* package, where an edge-betweenness algorithm selects the optimal topology, i.e. with high intra-community but low inter-community connections. The exon communities identified in the optimal topology correspond to the exon orthogroups returned by *ExOrthist*. Finally, *ExOrthist* offers the additional functionality to generate orthogroups restricted to two species when species pairwise gene orthopairs are provided as extra input.

### *ExOrthist* calibration and benchmarking

*ExOrthist* allows the user to differentially specify a set of conservation cut-offs across three evolutionary distance ranges (short, medium, long): (a) minimum global protein similarity (“prot_sim”); (b) minimum number of conserved intron positions (0, 1 or 2; “ int_num”); (c) minimum exon sequence similarity (“ex_seq”); and (d) minimum exon length ratio (shortest_exon/longest_exon; “ex_len”). This flexibility in the definition of cut-offs is particularly relevant, since the use of fixed conservation parameters across different evolutionary distances would result in noisy homology calls for closely related species and incomplete homolog detections for more distant species.

Minimum global protein similarity, or cut-off (a), is the minimum sequence similarity over the entire pairwise protein alignment for a pair of protein isoforms to be considered for further comparisons. While it has no effect on the downstream analyses, it avoids processing spurious gene orthologs or poorly annotated isoforms. The minimum number of conserved introns, or cut-off (b), refers to the requirement for both, one or none of the neighboring intron positions to be conserved, and hence inferred to be ancestral (Fig. 1A). A pair of introns from two genes are considered to be in conserved positions if they have the same phase (i.e. interrupt a codon in the same nucleotide position; Fig. 1A) and are in a comparable location of the protein alignment. In particular, the maximum distance allowed between the two conserved intron positions in the protein alignment depends on the overall protein similarity and percentage of unaligned residues (see Methods) [30]. In this study, we consider homologous internal exons only those for which both intron positions are conserved; therefore, this cut-off is always set to 2 in all the following analyses. However, by lowering the number of required conserved introns to one, internal exons sharing a single intron position can also be accommodated in the final orthologies.

Moreover, *ExOrthist* requires a minimum exon protein sequence similarity, or cut-off (c), for both the query exon as well as the neighboring exons, and a minimum exon length ratio, or cut-off (d), between the query exon and its match. Since these two cut-offs are the main determinants of exon homology calls, we aimed at defining the most optimal combination of cut-offs for each evolutionary distance range. For this purpose, we chose human and mouse, human and zebrafish and human and drosophila as representative species pairs for short, medium and long evolutionary distances, respectively. We first evaluated *ExOrthist*’s performance at a fixed exon length ratio (0.40) and with a minimum sequence similarity ranging from 0.1 to 0.9 (Fig. 2A-C). Here, it is expected that lower cut-offs will retrieve higher numbers of inferred homologous relationships. Therefore, we specifically aimed at identifying the highest sequence similarity cut-off yielding nearly maximal detection sensitivity. This means the cut-off at which the vast majority of potential exon homologs would be included in the final orthogroups but for which lower cut-offs would not substantially increase the number of detected conserved exons. In the case of short and medium evolutionary distance ranges, the number of conserved exons included in orthogroups reached saturation at sequence similarity 0.5 and 0.3, respectively, with lower cut-offs leading to only minor increases in the number of exons included in the orthogroups. Thus, we selected 0.3 and 0.5 as the default minimum sequence similarity cut-offs for short and medium evolutionary distance ranges. For the human -drosophila comparison, it was not possible to identify a saturation point at this long evolutionary distance, suggesting that a substantial fraction of exonic sequences has evolved beyond recognition. We therefore set the default value for minimum sequence similarity for long evolutionary distance ranges to 0.1.

**Fig. 2:**
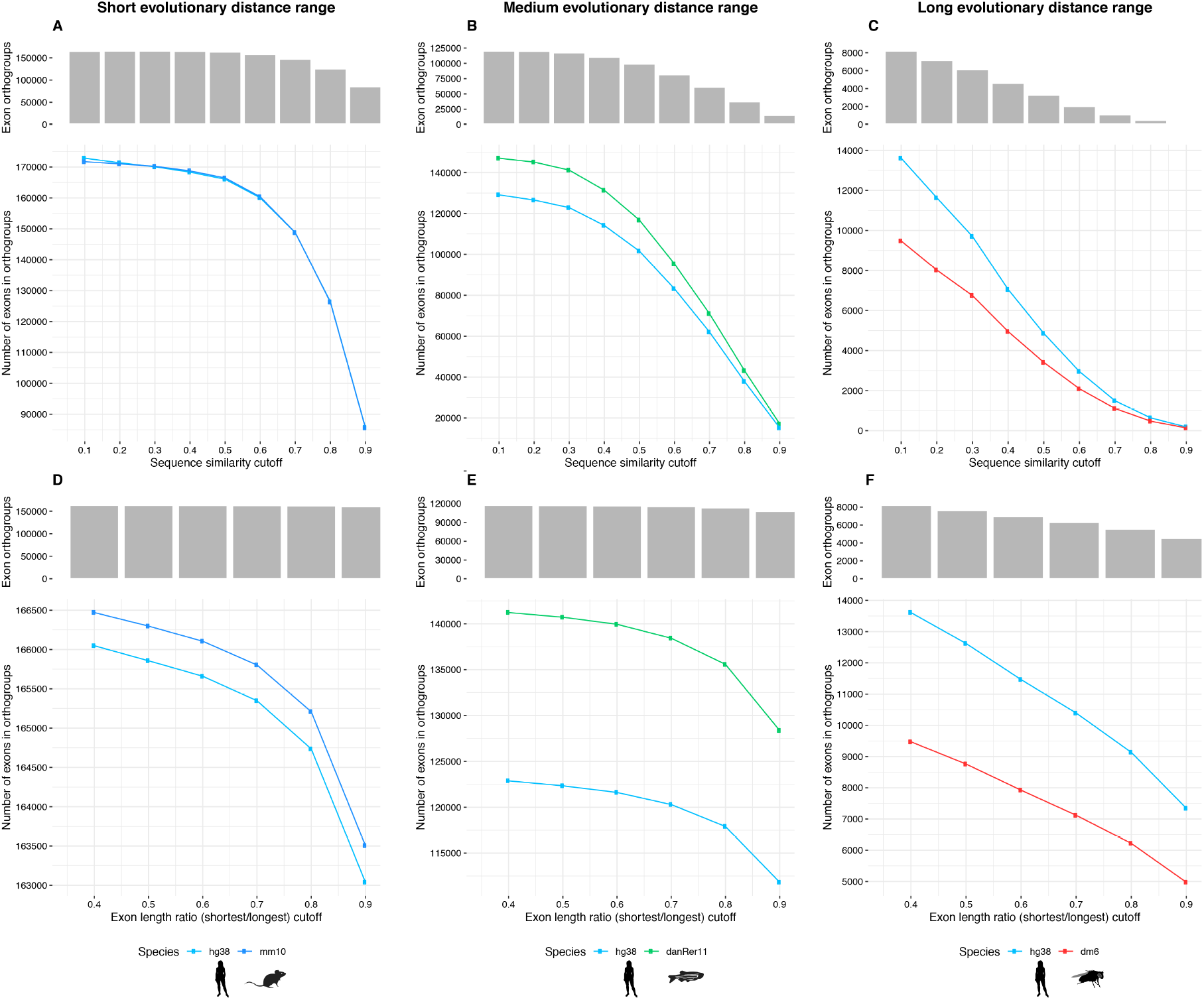
Calibration of *ExOrthist*’s conservation cut-offs. Estimation of default conservation cut-offs for each evolutionary distance range from two-species *ExOrthist* runs. **a**,**d**: human and mouse (short evolutionary distance range). **b**,**e**: human and zebrafish (medium evolutionary distance range). **c**,**f**: human and fruitfly (long evolutionary distance range). (**a-c**) The line charts depict the number of exons from each species included in *ExOrthist* orthogroups when different sequence similarity cut-offs are set [0.1-0.9]. The barplots on top represent the number of exon orthogroups retrieved by *ExOrthist* with the corresponding sequence similarity cut-off. All *ExOrthist* runs were performed with a fixed exon length ratio (shortest/longest) cut-off of 0.40. (**d-f**) The line charts depict the number of exons for each species included in *ExOrthist* orthogroups when different exon length ratio (shortest/longest) cut-offs are set [0.4-0.9]. The barplots on top represent the number of exon orthogroups retrieved by *ExOrthist* with the corresponding exon length ratio cut-off. *ExOrthist* runs were performed with the previously identified default sequence similarity cut-off (**d**: 0.5, **e**: 0.3, **f**: 0.1).

Next, using these default cut-offs for the minimum sequence similarity, we evaluated the effect of varying the minimum exon length ratio from 0.4 to 0.9 for each evolutionary distance range. As above, the number of inferred exon homologs and orthogroups saturated at an intermediate value for short and medium, but not for long, evolutionary distance ranges (Fig 2D-F). Therefore, we set the default values for minimum exon length ratio to 0.6 for short and medium and 0.4 for long evolutionary distance ranges, respectively. Data on the computing performance of *ExOrthist* for each type of run (short, medium, long) is provided in Supplementary Fig. 1 and in the respective *Nextflow* reports (Supplementary Files 1-6). As an example, an analysis involving two vertebrate species with many annotated isoforms (e.g. human and zebrafish, with ∼93K and ∼46K protein-coding transcripts, respectively), takes ∼2 hours of run time in a CPU cluster, requiring ∼70 hours of CPU time, 4.5GB of RAM memory and less than 10GB of disk space. Providing precomputed IPA alignments from previous *ExOrthist* runs (using the option *--prevaln*) reduces the necessary CPU time to less than 10 hours, saving more than 85% of the required computation time. We also implemented a configuration file for running the whole analysis on the Amazon cloud. For this test case, we uploaded input data into a S3 Bucket and used an AWS Batch spot queue with a bid percentage of 50% using optimal instance type nodes. The test run took around ∼13h to complete, with 40 CPU hours and a total cost below $10. Thus, there is no need for having a powerful HPC for running an analysis with well-annotated vertebrate species with *ExOrthist*.

We also aimed at assessing the sensitivity and accuracy of *ExOrthist*’s homology calls. Although there is currently no available software to infer exon homologies, we compared the output of *ExOrthist* with that obtained using a *liftOver*-based approach ([31] and see Methods). We generated exon orthogroups for all annotated exons in human, mouse and cow using *ExOrthist* with default cut-offs for short evolutionary distance. *ExOrthist* retrieved 148,255 1:1:1 exon triplets (83.3% of all exon orthogroups) and 25,832 (14.5%) orthogroups with a missing species (40.1% of which corresponded to cow) (Fig. 3A). Internal coding human exons showed higher levels of genomic conservation (i.e. had at least an ortholog in another species) than first and last exons (Fig. 3B), and, as expected, these conservation levels were strongly dependent on the average inclusion of the exon, with >99% and <18% of constitutive and cryptic internal exons having an ortholog in the other species, respectively (Fig. 3B). Importantly, the comparison with the *liftOver*-based pipeline showed a high degree of overlap (>97%) among the exons recovered in the orthogroups by each approach (Fig. 3C), with discrepancies largely corresponding to differences in annotations of exon variants between species. Moreover, the concordance between the orthology calls was ∼100% (i.e. a pair of exons from species1 and species2 was considered an orthopair by both methods) (Fig. 3C). Finally, we selected a well-studied gene orthogroup (*MBNL2*) to exemplify the output of the *exint_plotter* module (Fig. 3D).

**Fig. 3:**
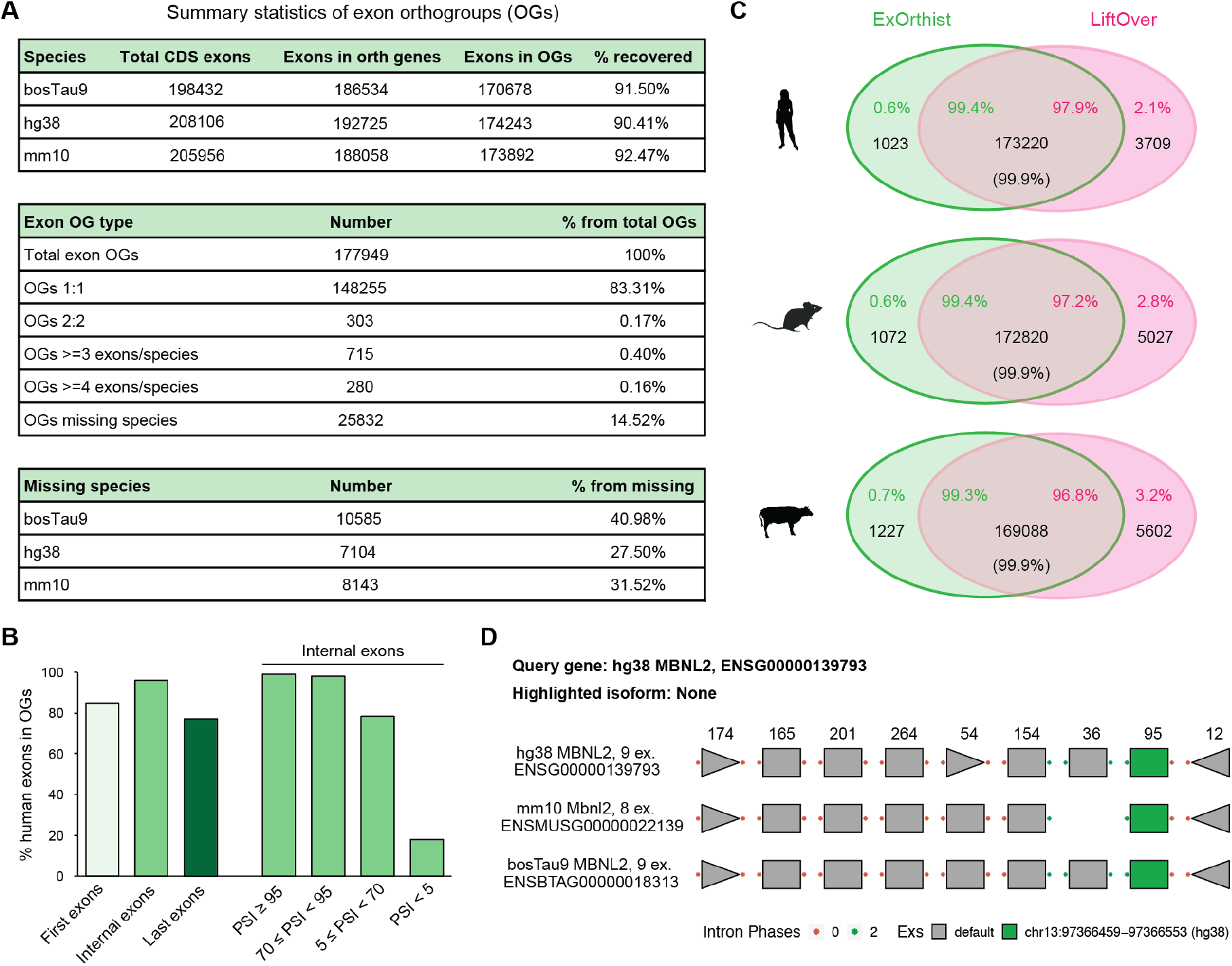
*ExOrthist* output and benchmarking. (**a**) Statistics of the exon orthogroups (OG) from an *ExOrthist main* run on three mammalian species (human, mouse, cow) as generated by *get_cluster_stats*.*pl*. (**b**) Percentages of conserved human coding exons based on their location within the gene (first, internal, last) or on their average inclusion level (for internal exons only), as obtained from *VastDB* [27]. (**c**) Venn diagrams showing the number of exons (black) included in either *ExOrthist* orthogroups (green), *liftOver*-based orthogroups (pink) or both (intersection) for human, mouse and cow. The percentages of shared and not-shared exons between *ExOrthist* and *liftOver*-based orthogroups are reported in the corresponding area. The percentage in brackets refers to the concordance between the 1:1 exon pairs in the target species (e.g. mouse and cow for human exons) retrieved by both methods. (**d)** Plot of the exon-intron structure of human *MBNL2* and its mouse and cow orthologs as generated by the *exint_plotter* module. Exons in the different species falling in the same exon clusters are vertically aligned, and a randomly chosen exon of interest is highlighted in green.

### Using *ExOrthist* to investigate asymmetric exon evolution after gene duplication

One important feature of *ExOrthist* is that it affords exon homology inferences not only for cases of clear 1:1 orthologs but also in scenarios with different levels of paralogy. Therefore, to assess this functionality, we performed an exon comparison between *Xenopus tropicalis* and the allotetraploid *Xenopus laevis*.Since the whole genome duplication of *X. laevis* is relatively recent (17-18 million years ago), most of its genes can be readily assigned to any of the two ancestral subgenomes (named L and S, respectively; Fig 4A) [32]. Although both ancestral copies (homeologs) have been retained for multiple genes, a substantial fraction has asymmetrically retained a single copy, and this has most often corresponded to the one in the L subgenome (Fig. 4B)[32]. To investigate exon evolution after whole genome duplication, we used *ExOrthist* to generate exon orthogroups within a pre-defined set of 8,806 *X. leavis* homeolog pairs and their respective *X. tropicalis* single orthologs (1:2 gene orthologs) [32]. *ExOrthist* identified 69,035 (88.0%) 1:2 exon orthologs as well as 9,377 (12.0%) 1:1 relationships, which indicate lack of exon homology for one gene homeolog in *X. laevis* after genome duplication (Fig. 4B). Interestingly, similar to the pattern of retention of gene duplicates, we found that gene copies from the L sub-genome more often retained ancestral exon homology compared to the S sub-genome (Fig. 4B; 57.7%, *P* = 7.8e-48, two-sided Binomial test).

**Fig. 4:**
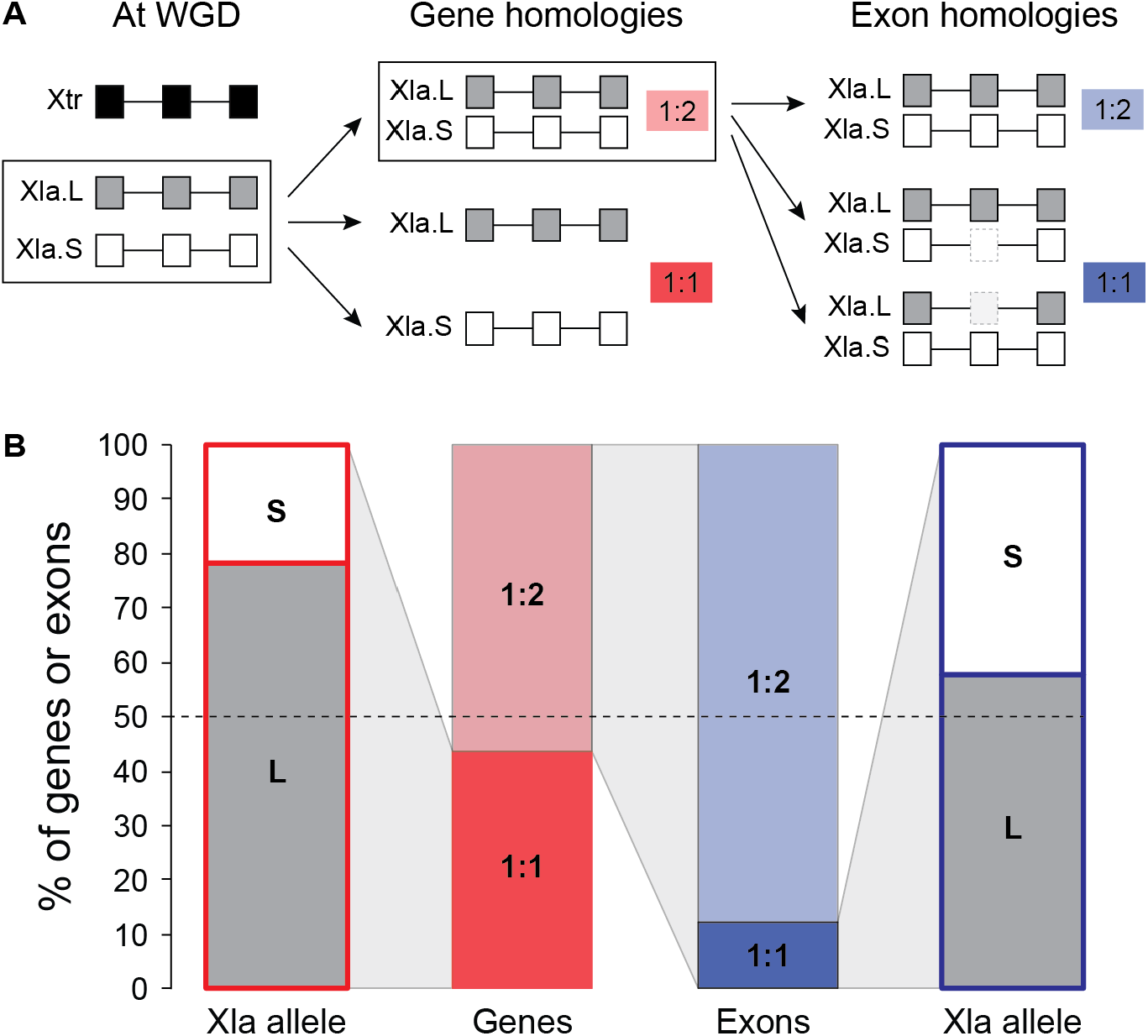
*ExOrthist* paralogy inference in the *Xenopus* genus. (**a**) Scheme depicting the most common fates for the *X. laevis* (Xla) paralogs (genes and exons) generated by a whole genome duplication (WGD) occurred well after the split between Xla and *X. tropicalis* (Xtr). Each copy of a paralogous gene pair belongs to a different subgenome (S or L). Both gene paralogs can be conserved, giving rise to 1:2 orthologs (Xtr:Xla). Alternatively, one gene copy can be lost (either from the S or the L subgenome), re-establishing a 1:1 orthology relationship. The same classification is applied to paralogous exons in 1:2 gene orthologs. Ancestral exons may be conserved in both genes copies (1:2) or lost by either the S or L copy (1:1). (**b**) Percentages of 1:1 and 1:2 genes (red) or exons (blue) in the Xla genome, with the relative percentages of S and L alleles for the 1:1 genes/exons. Cases for which the L or S origin was not defined were excluded from the plot.

### Evaluation of genomic and regulatory conservation of alternatively spliced exon sets

A major application of *ExOrthist* is the assessment of the evolutionary conservation of alternatively spliced exon networks (also referred to as exon programs). These networks consist of sets of exons that are regulated in a coordinated manner in a specific tissue (e.g. brain, muscle), usually through the action of tissue-enriched alternative splicing factors, and are known to play crucial roles in cellular differentiation and physiology [33, 34]. Moreover, some sets of exons have been shown to be jointly mis-regulated in human pathologies, including cancer or mental disorders [35, 36], and the evolutionary comparison between human and model organisms is a powerful approach to identify potential pathogenic targets [37]. The *ExOrthist compare_exon_sets* module allows the evaluation of the evolutionary conservation of exon sets between two species from a genomic and regulatory perspective, both at the gene and exon level (Fig. 5A). At the gene level, *ExOrthist* assesses the proportion of exons in genes that have orthologs in the target species and the proportion of those orthologs that also harbor regulated exons. Similarly, exons are individually evaluated to infer what proportion have orthologs in the other species (referred to as “Genome-conservation” [38]) and for what proportion those orthologs are also regulated (“Regulation-conservation” [38]). Additionally, for all gene orthologs containing regulated exons in both species, the *compare_exon_sets* module performs a pairwise comparison to assess whether the exon pairs are: (i) orthologs, (ii) best-hits (i.e. they are considered the best matching exon by the *main* module, but do not meet all the requirements to be considered proper homologs), non-orthologs, or (iv) unclear. In particular, *ExOrthist* relies on several aspects to confidently assign two exons as non-orthologs (Supplementary Fig. 2), allowing the inference of evolutionarily independent alternative splicing patterns.

**Fig. 5:**
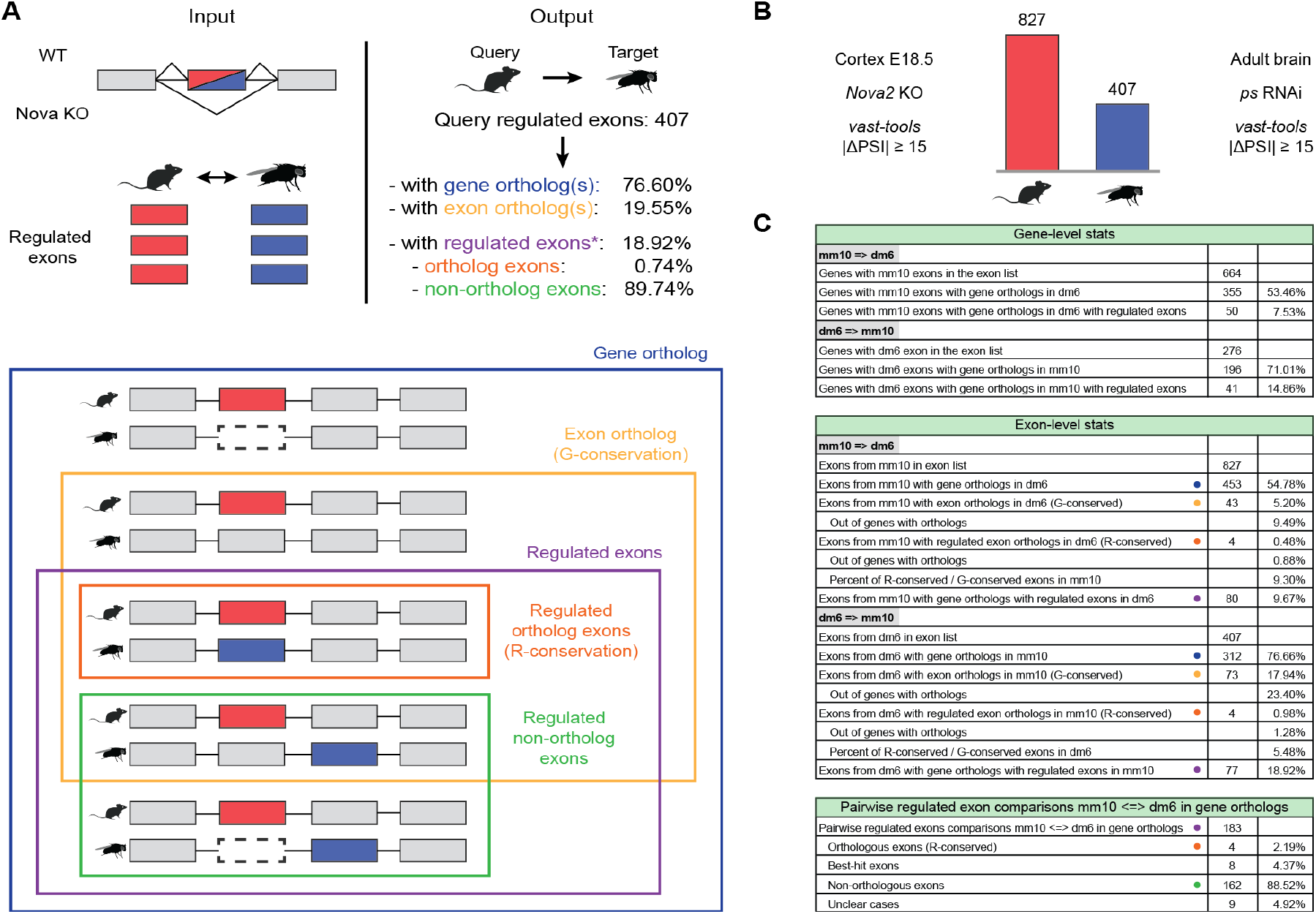
*ExOrthist compare_exon_sets* module. (**a**) Schematic representation of the inputs required by the *compare_exon_sets* module and of the provided output. For a given query regulated exon (red): “Gene ortholog”, the query exon belongs to a gene that has orthologs in the target species; “Exon ortholog”, the target exon has at least one valid exon ortholog as defined by *ExOrthist* in the target species (referred to as Genome-conservation or G- conservation); “Regulated exons”, both the query exon and a target exon (blue) are regulated, whether they are orthologs (“Regulated exon orthologs (R-conservation)”) or not (“Regulated non-ortholog exons”). (**b**) Number of *Nova*-dependent exons in mouse and fruitfly as identified by *vast-tools* (*ps*: *pasilla*, drosophila homolog of mouse *Nova1* and *Nova2*). (**c**) Output of the *compare_exons_sets* module when run on mouse and fruitfly *Nova*-regulated exon sets presented in (b). Color dots correspond to scenarios depicted in (a).

We used *ExOrthist* to compare sets of exons regulated by the splicing factor *Nova* in mouse and fruitfly. For this purpose, we ran *vast-tools* [27] to obtain changes in inclusion levels (ΔPSIs) for all exons upon *Nova2* depletion in mouse embryonic cortex [39] and upon knockdown of the fruifly ortholog (*ps*) in whole adult brains [40]; we then defined as *Nova*-dependent exons in each species those with a |ΔPSI| > 15 (827 in mouse and 407 in fruifly, respectively; Fig. 5B). In parallel, we ran *ExOrthist main* for mouse and fruitfly with default cut-offs for the long evolutionary distance range and providing all exons evaluated in *vast-tools* as additional exons for each species through the *--extraexons* option. We then utilized the *compare_exon_sets* module to compare the mouse and fruitfly *Nova*-dependent exon sets and identify conserved exons. Despite the expected low conservation [41, 42], *ExOrthist* identified 43 (5.2%) mouse *Nova*-dependent exons sharing orthogroups with fruitfly exons, 4 of which were also impacted by *Nova* depletion in the insect species (Supplementary Table 1). However, the most remarkable finding of this investigation was evidence for shared *Nova* regulation of orthologous genes through regulation of non-orthologous exons: 80 mouse exons (in 50 genes) were in gene orthologs harboring *Nova*-dependent exons in fruitfly (77 exons in 41 genes; Fig. 5C), the vast majority of which (162/183 pairwise comparisons, 88.5%) corresponded to non-orthologous exon pairs in the two species, compared to the 4 (2.2%) cases with regulatory conservation and 8 (4.4%) additional pairs that were best exon hits although they did not pass all the homology filters (Supplementary Table 2). These results highlight a major evolutionary convergence of *Nova*-regulated exon networks impacting orthologous genes, as seen also for other splicing factor networks [42–44].

## Discussion

Exon gains and losses have the potential to drive diversification of gene structure and function across eukaryotes. Moreover, alternative splicing of specific exons can greatly expand transcriptome and proteome diversity, especially in metazoans. In this context, evolutionary conservation both at the genomic and the regulatory level of alternative splicing has been widely regarded as a proxy for functional relevance [38]. A key step to assess such conservation is the inference of exon homology relationships among species. However, while multiple tools have been developed for gene homology inference, there is currently no available software to derive exon homology relationships. Here, we have introduced *ExOrthist*, the first software specifically designed to infer exon orthology and paralogy relationships taking into account the unique characteristics of exon and intron evolution across different evolutionary distances. Therefore, unlike gene-oriented software, *ExOrthist* does not rely on comparisons of exon sequence similarity in isolation, but also makes use of the full intragenic context. For this reason, *ExOrthist* identifies potential exon homologs through global protein alignments, and explicitly requires conservation of the neighboring intron positions as well as of individual exon sequences within these global alignments.

Evolution of intron positions has been extensively studied, revealing a generally low rate of intron gain and loss within most eukaryotic groups [45–48], despite some remarkable exceptions (e.g. [49, 50]). Comparison among lineages showed conservation of relatively high fractions of ancestral introns in various clades, as well as substantial remodeling of intron-exon architectures [47, 51]. Since, by definition, intron insertions define exons, conservation of ancestral intron positions is a requirement for true exon homology. Thus, it is expected that a large fraction of exons is conserved within vertebrates (Fig. 3, [5, 52, 53]), but not between species with largely divergent intron-exon architectures (e.g. between mammals and dipterans; Fig. 5). That is, even if the compared coding sequences are orthologous, the actual exon entities encoding them may not be. To our knowledge, *ExOrthist* is the first framework that incorporates the conservation of intron positions and other features of the gene context to draw the fundamental distinction between conserved exon sequences and conserved exon entities. In order to clearly distinguish between these two scenarios, *ExOrthist* also provides the best hit for each exon within each target gene homolog, even when it does not meet the necessary requirements to be considered an exon homolog.

Since *ExOrthist* allows users to work at any evolutionary distance and with species for which only a genome sequence and a standard gene annotation is available, our software can be used for both model and non-model organisms. For this reason, we envision that *ExOrthist* will enable unprecedented evolutionary studies, both in terms of species representation and biological questions, opening multiple new venues of research on genome evolution and alternative splicing. To name a few, these could include the identification of deeply conserved alternative splicing events, the study of the origin and evolution of tissue-specific exon networks, the investigation of intragenic evolutionary patterns, the inference of phylogenies based on *bona fide* exon orthologs, and the construction of large multi-species databases interconnected not only through gene homology relationships (e.g. *Ensembl*), but also through exon homology relationships (e.g. *VastDB* [27]).

## Conclusions

*ExOrthist* is the first tool specifically designed to infer exon orthogroups and orthopairs across any evolutionary distance. It also contains additional modules to visualize conservation of intron-exon architectures and to perform evolutionary comparisons of exon sets of interest. *ExOrthist* will thus facilitate research on genome evolution at the intragenic level and the identification of highly conserved, and thus more likely functional, alternatively spliced exons.

## Methods

### Algorithm of *ExOrthist main* module

The *ExOrthist main* module requires a minimum set of inputs (annotation files, genome files, evolutionary distance information and gene orthogroups) to return the complete set of exon orthopairs and orthogroups for a chosen group of species. The following sections (A, B, C, D) describe the algorithm applied by *ExOrthist*, as well as the extra steps implemented when additional inputs are provided. The module relies on software and libraries stored within a docker image whose recipe is bundled together with the rest of the pipeline. The image is available at docker hub (https://hub.docker.com/r/biocorecrg/exon_intron_pipe) and is automatically retrieved by *ExOrthist* and eventually converted into a singularity image if the option *-with-singularity* is specified. To run the program, a user only needs to install *Nextflow* and one Linux container, either *Docker* or *Singularity*. Further details can be obtained in the README section at https://github.com/biocorecrg/ExOrthist.

#### A. Input generation

After checking for the presence of all required inputs and the consistency between gene orthogroups and the annotated gene IDs in the GTF files, *ExOrthist* generates files with annotation information for each considered species, which will be the input of all species pairwise comparisons (see online README for a detailed list of all the files). Although, by default, only annotated exons are considered, *ExOrthist* can integrate non-annotated exons into its homology inferences by providing a list of gene identifiers plus exon coordinates (e.g. exons identified by *vast-tools* or any other software to detect and/or quantify alternative splicing). If non-annotated exons are provided, *ExOrthist* generates a new GTF file where a novel transcript for each of the non-annotated exons is introduced. *ExOrthist* first maps the upstream and downstream exons (provided in the input file) to the annotated transcripts. When both neighboring exons match the same transcript, this becomes the template of the novel transcript to which the non-annotated exon is added; otherwise, *ExOrthist* uses partial matches of the upstream or downstream exon to identify the template transcript and create the new annotation.

*ExOrthist* allows the user to specify different conservation cut-offs depending on the evolutionary distances between the compared species, which have to be declared in a specific input file. In particular, four evolutionary cut-offs can be fine-tuned for three evolutionary distance ranges (short, medium, long): (a) minimum global protein sequence similarity for a pairwise isoform alignment to be considered for the orthology call, (b) minimum number of conserved intron positions, (c) minimum sequence similarity required for orthologous exons, and (d) minimum ratio between the lengths of an exon pair (length of the shortest/longest exon). Cut-offs (b), (c) and (d) will be utilized in the later steps of the pipeline to filter out non-homologous matches from the pool of candidate exon orthopairs for each pair of species (see section C).

#### B. Parsing of IPA alignments

*ExOrthist* then works by species pairs and within gene orthogroups. For each species pair and gene orthogroup, it first generates pairwise IPA alignments between all non-redundant protein isoforms of species1 vs. all non-redundant protein isoforms of species2 (Fig. 1C). Considering species1 as query and species2 as target (and vice versa), *ExOrthist* parses the IPA alignments to derive:

- Protein conservation information: species-wise percentages of sequence identity, sequence similarity, and gaps between the two proteins are calculated. Alignments are not considered for further processing unless the global sequence similarity (cut-off (a)) reaches the minimum protein similarity specified for the relative evolutionary distance (see section A) for both species, or twice (2x) the minimum protein similarity cut-off for one of the species.
- Exon conservation information: for each query exon, *ExOrthist* selects all exon matches in all target isoforms. In case of a query exon matching multiple target exons in the same isoform each covering ≥15% of the query exon sequence, each exon pair is specifically realigned and the best match is selected based on sequence similarity.
- Intron conservation information: *ExOrthist* tries to match each query intron with a target intron in the IPA alignment. Depending on the protein conservation, the width of the searching window around the query intron changes:
  - if % similarity ≤ 30 or % gaps ≥ 30: width = 10
  - if % similarity ≥ 30 and % similarity < 50: width = 8
  - if % similarity ≥ 50 and % similarity < 70: width = 6
  - if % similarity ≥ 70 and % similarity < 80: width = 4
  - if % similarity ≥ 80 and % similarity < 90: width = 3
  - if % similarity ≥ 90: width = 2

This window is further increased by the average number of gaps (i.e. “-”) within ten positions at each side of the alignment. Then, if a target intron is not found within the search window, conservation of the query intron is set to zero. If a match is found and the phases of the two introns are equal, intron conservation is rated 10. If the phases of the two introns are different, intron conservation is rated −10. Next, if the two intron positions are not perfectly aligned, the number of residues by which they are separated in the alignment is subtracted to the intron conservation value. Thus, the maximum intron conservation value is equal to 10 for a perfectly aligned pair of introns in the same phase, and decreases progressively to indicate no intron conservation (0) or presence of non-conserved introns with different phase (negative values). The intron position conservation value is later translated into a partial score used to evaluate pairwise exon orthology relationships.

*ExOrthist* efficiently parallelizes all alignments and realignments thanks to the *Nextflow*’s management system, and later joins all the best isoform-wise matches for a given species pair. If the output of a previous *ExOrthist* run is provided (*--prevaln* option), the alignment and realignment steps are skipped for all the query and target isoform pairs that have already been aligned.

#### C. Scoring and best match selection

*ExOrthist* keeps executing the processes separately for each species pair. In here, it selects the best match for each exon in the query species among all the isoform matches of each homologous gene in the target species. The selection is based on five partial scores reflecting the conservation of the previously assessed features of the exon-intron context (section B):

- **(1)-(2):** Conservation of the (1) upstream and (2) downstream intron phases: ranging [-0.25, 0.25], obtained from multiplying the intron conservation value from B by 0.025. Positive values indicate phase conservation, while deviations from the maximum score reflect shifts in the intron positions within the IPA.
- **(3):** Conservation of the exon sequence: ranging [0, 0.2]. 0.2 indicates 100% sequence similarity.
- **(4) - (5):** Conservation of the (4) upstream and (5) downstream exon sequences: ranging from [0, 0.15]. 0.15 indicates 100% sequence similarity.

These partial scores for internal exons are then summed up to generate a global score ranging [0,1]. In the case of first or last exons, for which scores (2) and (5) or (1) and (4) do not exist, respectively, the global score is divided by 0.6 to make it range [0,1]. For each species pair, different filtering criteria will then be applied based on their evolutionary distance and the specified conservation cut-offs. The up/downstream intron position conservation, the exon sequence conservation and the up/downstream exon sequence conservation are sequentially evaluated in the filtering.

- (**1**) and/or **(2**), depending on cut-off (a), are required to be positive in all the target-gene best matches (i.e. the intron positions need to be in the same phase and within the allowed window, even if not necessarily in the same location). If the intron phase(s) are conserved, sequence conservation is evaluated. For first and last exons, only (2) and (1) are evaluated, respectively.
- (**3**) is required to be ≥ *min_ex_sim*\*0.2, where *min_ex_sim* is the minimum sequence similarity specified for species1-species2 evolutionary distance range (cut-off (b)).
- (**4**) and **(5)** are required to be ≥ *min_ex_sim*\*0.15, where *min_ex_sim* is the minimum sequence similarity specified for species1-species2 evolutionary distance range (cut-off (b)). For first and last exons, only (5) and (4) are evaluated, respectively.
- Extra filter: the length ratio between two compared exons (shortest/longest) should not be lower than the minimum exon length ratio specified for species1-species2 evolutionary distance range (cut-off (c)).

In the end, among these pre-filtered matches, the match with the highest global score is selected for each query exon and target gene. If no target exon passes all cut-offs, the one with the highest global score is considered the best hit for the target gene. While the *ExOrthist*’s logic requires a query exon to match a unique target exon, each target exon can potentially be matched by more query exons. Overlapping variants of the query exons (i.e. alternative forms of the same exon due to alternative splice donor/acceptor sites) might be present in the pool of filtered matches. In order to univocally derive orthologous relationships for each exon, *ExOrthist* selects the form with the highest number of matches (among all target genes) as representative of its overlap group. In addition, *ExOrthist* accepts custom exon orthopairs (from *liftOver* or manual curation) to be considered in the final exon orthology inference through the *--bonafide_pairs* option. For this purpose, *ExOrthist* incorporates a custom script (*get_liftovers*.*pl*) to allow users to retrieve *liftOver*-based exon orthopairs for species pairs with available *liftOver* files. If provided, these exon orthopairs will be directly integrated in the generation of the exon orthogroups graphs (next section).

#### D. Exon clustering

After deriving exon homologous relationships between each pair of species, *ExOrthist* works on the combined orthopair information to infer the exon orthogroups. For each gene orthogroup, *ExOrthist* builds a directed graph (through the R *igraph* package v1.2.6 [54]) with exons as nodes and their pairwise homology relationships represented as edges. In case of a best reciprocal match between homologous exons, two directed edges will be drawn between the correspondent nodes. *ExOrthist* then applies the R *igraph* edge-betweenness algorithm to select the optimal graph topology, with communities highly intra-connected and lowly inter-connected. Although the directionality of the graph is not considered by the edge-betweenness algorithm, the reciprocity of the matches is represented by the number of edges. The exon communities identified in the optimal topology correspond to the multi-species exon orthogroups returned by the pipeline. For each exon, *ExOrthist* also computes a Membership Score (MS), which reflects its degree of similarity to all the exons belonging to the same orthogroup (OG). The MS is defined as follows:

MS = (IN_degree + OUT_degree + N_reciprocals) / (2*(TOT_exons_in_OG - SPECIES_exons_in_OG) + (TOT_genes_in_OG - SPECIES_genes_in_exOG)) with:

- IN_degree: number of exon matches from the other exons in the orthogroup (i.e. the considered exon is target).
- OUT_degree: number of exon matches to the other exons in the orthogroup (i.e. the considered exon is query).
- N_reciprocal: number of reciprocal matches (i.e. query exon is a match of its target exon and vice versa).
- TOT_exons_in_OG: number of exons in the orthogroup.
- SPECIES_exons_in_OG: number of exons from the same species present in the exon orthogroup.
- TOT_genes_in_OG: total number of genes in the original gene orthogroup.
- SPECIES_genes_in_exOG: number of genes from the same species that contribute with exons to the exon orthogroup.

### Calibration of *ExOrthist*’s default conservation cut-offs

We first selected three pairs of species as representatives of each evolutionary distance range. Human-mouse for short (hg38-mm10; estimated divergence: 90 million years ago [MYA] [55]), human-zebrafish for medium (hg38-danRer11; estimated divergence: 435 MYA) and human-fruitfly for long (hg38-dm6; estimated divergence: 797 MYA). We then used *Broccoli* v1.0 with default settings [18] to infer each pairwise gene orthogroups. We removed all the gene orthogroups containing more than 20 genes, selecting 16,034 orthogroups for hg38-mm10, 11,571 for hg38-danRer11 and 5,142 for hg38-dm6.

In order to identify the optimal minimum exon sequence similarity (parameter (c)) and minimum exon length ratio (shortest/longest) (parameter (d)), we systematically analyzed the *ExOrthist* performance with different combinations of those parameters. First, we kept the minimum length ratio fixed at 0.4 and we performed nine *ExOrthist* runs for each species pair starting from a minimum sequence similarity of 0.1 and increasing it by 0.1 in each of the following runs up to 0.9. We then selected the optimal sequence similarity cut-off for the short and medium evolutionary distance ranges depending on when saturation in the number of conserved exons was reached (0.5 and 0.3, respectively) (Fig. 2A-C). For long evolutionary distances, we selected 0.1 as default sequence similarity cut-off, since saturation was not reached.

Second, we kept the minimum sequence similarity fixed at the identified default parameter for each evolutionary distance range, and we performed five *ExOrthist* runs for each species pair starting from a minimum exon length ratio of 0.4 and increasing it by 0.1 in each of the following runs up to 0.9. The saturation in the number of conserved exons was reached at an exon length ratio cut-off of 0.6 for both short and medium evolutionary distance ranges. For long evolutionary distances, we selected 0.4 as the default exon length ratio cut-off, since saturation was not reached (Fig. 2D-F). All the mentioned runs were executed with minimum protein similarity=0.25 (short), 0.20 (medium), 0.15 (long) and fixed minimum number of conserved introns=2.

### Comparison of *ExOrthist* and *liftOver* performances

To benchmark *ExOrthist* at a short evolutionary distance range, we implemented a *liftOver*-based approach following previous studies [27, 31]. Briefly, we downloaded the *liftOver* chain alignment files for each pairwise comparison (among human, mouse and cow; hg38, mm10 and bosTau9) and performed the *liftOver* for all annotated exons from each species against the other two. As recommended by the developers, we used the following parameters: -minMatch=0.10 -multiple -minChainT=200 -minChainQ=200, as implemented in the script *get_liftovers*.*pl* provided as part of *ExOrthist*. Then, to make the comparisons between *ExOrthist* and *liftOver* fairer, we did not require the *liftOver* matches to have any canonical dinucleotide, and we considered only lifted coordinates that shared a junction with an annotated exon in the target species and in the same gene orthogroups used to run *ExOrthist*. From these exon pairs, we generated exon orthogroups by running the final scripts of the *ExOrthist main* module (*D2_cluster_EXs*.*R* and *D3_format_EX_clusters_output*.*pl*).

To obtain *ExOrthist* exon orthogroups, we used *Broccoli* v1.0 [18] with default settings to infer gene orthogroups for human, mouse and cow. We filter out 53 orthogroups containing more than 20 genes, restricting the following analysis to 17,594 orthogroups. We ran *ExOrthist* with conservation cut-offs set at the default values for short distance species (minimum sequence similarity=0.5; minimum exon length ratio=0.6, minimum global protein similarity=0.25 and minimum number of conserved introns=2).

To compare the exon orthogroups from both approaches, we performed two analyses. First, we asked, for each species and for each approach, how many annotated exons were found in orthogroups with exons from each of the other two species and plotted the overlapping fraction (Fig. 3C). Since exons may have alternative splice acceptor and/or donor sites, we used *bedtools intersect* to compute the overlap between the two orthogroup files. Second, for exons with a 1:1 match in another species in both exon orthogroup files, we assessed the concordance of these orthology assignments (i.e. whether an exon in species1 was paired to the same exon in species2 by both approaches), obtaining virtually 100% concordance.

### Investigation of exon evolution after whole genome duplication

We obtained the 1:2 and 1:1 gene orthogroups between *X. tropicalis* (v9.0) and *X. laevis* (v9.1) from [32] and the respective genome annotations from Xenbase. Genome annotations were downloaded as GFF3 files and converted to GTF format using *gffread* [56] and custom scripts. We generated exon orthogroups by running *ExOrthist* with default conservation cut-offs for short evolutionary distance (minimum global protein similarity=0.25, number of required introns=2, minimum sequence similarity=0.5; minimum length ratio=0.6), and we subsequently quantified the exon orthogroups (1:2 and 1:1) generated from the 8,806 1:2 gene orthogroups. Information about the *X. laevis* subgenome (L or S) was obtained from [32], and was used to parse the 1:1 gene and exon orthogroups for the corresponding missing orthologs. We considered an exon to have 1:1 or 1:2 homologs only based on the exon orthogroups, and did not take the best hit information into account.

### Comparison of *Nova*-dependent exon sets

We used *Broccoli* v1.0 [18] with default parameters to infer mouse-fruitfly (mm10-dm6) gene orthogroups. We selected all the orthogroups (5,207) containing less than 20 genes. We ran *ExOrthist* with default conservation cut-offs for long distance evolutionary range (minimum global protein similarity=0.15, number of required introns=2, minimum sequence similarity=0.1; minimum length ratio=0.4), adding non-annotated exons identified by *vast-tools* [27]. To identify *Nova*-dependent exons, we run *vast-tools* v2.5.1 for two datasets characterized by deficiency of Nova orthologs in each species. In particular, we used an experiment with *Nova2* depletion in mouse embryonic cortex at the E18.5 stage [39] and another one with downregulation of *pasilla* (*ps*) in adult fly whole brains [40]. To obtain inclusion levels for all exons, we employed *vast-tools align* and *combine* with default parameters for mm10 and dm6 (VASTDB libraries: vastdb.mm2.23.06.20.tar.gz and vastdb.dme.23.06.20.tar.gz) [27]. All replicates for control and depleted samples were merged into a single sample using *vast-tools merge* to increase read depth, and *vast-tools compare* with default parameters (minimum change in inclusion levels [min_dPSI] of 15) was employed to define *Nova*-dependent exons (827 exons in mouse and 407 in fruitfly, respectively; Fig. 5B). Evolutionary comparison of both sets was done using the *ExOrthist compare_exon_sets* module with default options and providing two exon sets.

## Code availability

The *ExOrthist* is an open software publicly available on Github (https://github.com/biocorecrg/ExOrthist) under a MIT license.

## Authors’ contributions

YM and MI conceived the software. YM developed the core of the *main* module, with contributions from FM, MI, DB and SWR. FM developed the *exint_plotter* module. MI wrote the *compare_exon_sets* module with input from FM. LC, FM and JP made the *Nextflow* implementation. YM, FM and MI performed computational analyses. FM and MI wrote the manuscript and made the figures, with input from the other authors. All the authors critically reviewed and approved the final manuscript.

## Acknowledgments

Animal silhouettes were obtained from http://phylopic.org.

## Supplementary Table Legends

**Supplementary Table 1 - Orthogroups output from *compare_exon_sets***. List of orthogroups for the regulated exons provided as input for *compare_exon_sets*. INFO column corresponds to the regulatory information provided.

**Supplementary Table 2 - Pairwise comparison output from *compare_exon_sets***. For all regulated exons in orthologous genes in the two species, each pairwise comparison is reported. This includes information about exon and host gene, as well as the conservation call performed by *compare_exon_sets* (CONSERV_CALL). The terms correspond to those described in Supplementary Fig. 2.

## Supplementary Figures

**Supplementary Fig. 1.**
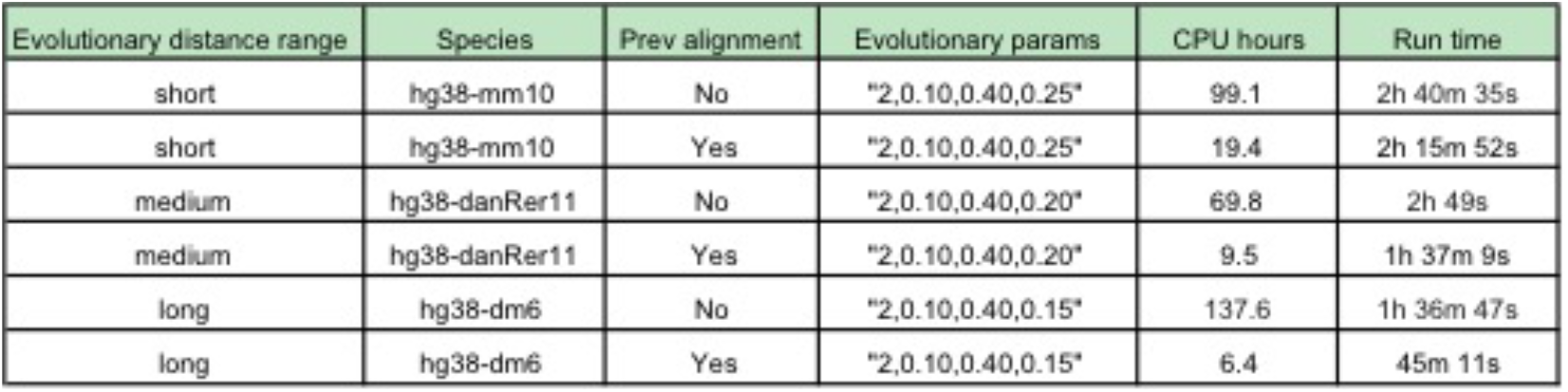
Performance of *ExOrthist*. CPU hours and run time is provided for each of the runs, as well as information about the runs (species, cut-offs, and whether or not the run included previous alignments). The cut-offs are in the format required for the *ExOrthist* run: “int_num,ex_seq,ex_len,prot_sim”. Additional information can be found in the Nextflow report for each of the runs (Supplementary Files 1-6).

**Supplementary Fig. 2.**
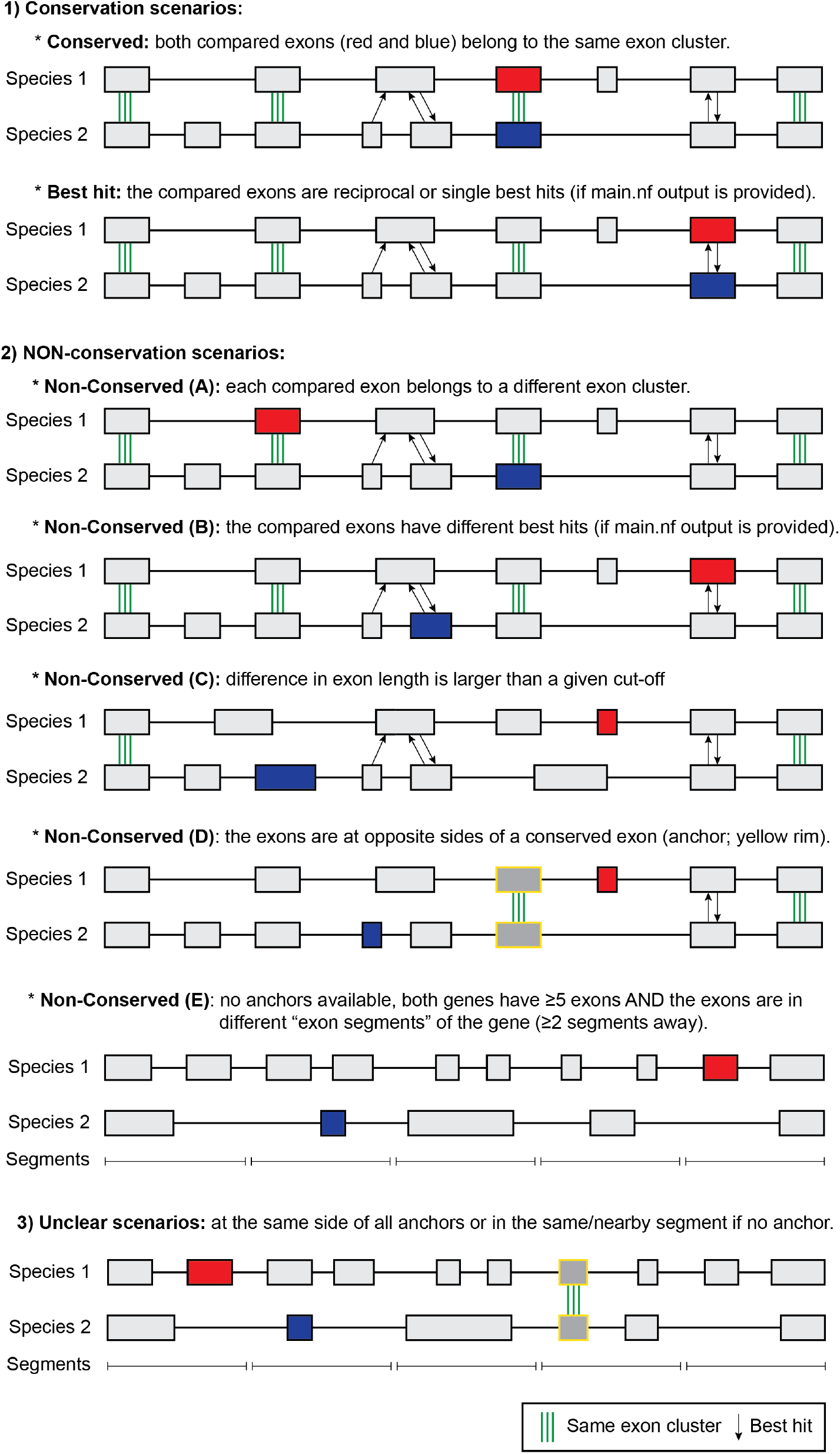
Scenarios of pairwise exon comparisons in orthologous genes by *compare_exon_sets*. Each non-conservation scenario is evaluated in a consecutive manner (i.e. if the conditions for A are met, the rest of scenarios are not assessed).

