## Supplementary Files 1-6 for "*ExOrthist*: a tool to infer exon orthologies at any evolutionary distance": Supplementary_File_2-hg38_mm10-0.10_0.40-withaln.html

[shrivelled\_faggin] Nextflow Workflow Report


Nextflow Report


- Summary
- Resources
- Tasks

[shrivelled\_faggin]

### Nextflow workflow report

#### `[shrivelled_faggin]`

Workflow execution completed successfully!

Run times
:   04-Jan-2021 14:03:48 - 04-Jan-2021 16:19:26
    (duration: **2h 15m 39s**)

15607 succeeded

0 cached

0 ignored

0 failed

Nextflow command
:   ```
    nextflow main.nf -with-report reports/hg38_mm10_withaln.html -with-trace
    ```

CPU-Hours
:   `19.4`

Launch directory
:   `/nfs/users/mirimia/fmantica/projects/regulatory_ancenstry/src/nextflow/EXORTHIST/ExOrthist`

Work directory
:   `/nfs/users/mirimia/fmantica/projects/regulatory_ancenstry/src/nextflow/EXORTHIST/ExOrthist/work`

Project directory
:   `/nfs/users/mirimia/fmantica/projects/regulatory_ancenstry/src/nextflow/EXORTHIST/ExOrthist`

Script name
:   `main.nf`

Script ID
:   `26c49de1d14c5feaed88006fc925ac34`

Workflow session
:   `83a68b08-c968-4a3e-bca6-eec58ee6063c`

Workflow profile
:   standard

Workflow container
:   `biocorecrg/exon_intron_pipe:0.2`

Container engine
:   `singularity`

Nextflow version
:   version 20.07.1, build 5412 (24-07-2020 15:18 UTC)

## Resource Usage

These plots give an overview of the distribution of resource usage for each process.

#### CPU

- Raw Usage
- % Allocated

#### Memory

- Physical (RAM)
- Virtual (RAM + Disk swap)
- % RAM Allocated

#### Job Duration

- Raw Usage
- % Allocated

#### I/O

- Read
- Write

## Tasks

This table shows information about each task in the workflow. Use the search box on the right
to filter rows for specific values. Clicking headers will sort the table by that value and
scrolling side to side will reveal more columns.

Values shown as:

Human readable
Raw values

(tasks table omitted because the dataset is too big)

Generated by Nextflow, version 20.07.1
